# Extracellular Vesicles Mediate Glucose Regulation by GLUT4-Overexpressing Engineered Muscle Tissue in T2D Mice

**DOI:** 10.1101/2025.07.08.663248

**Authors:** Hagit Shoyhet, Yahel Cohen, Yifat Herman Bachinsky, Irit Rosenhek-Goldian, Roman Kamyshinsky, Eran Horenstein, Shulamit Levenberg

## Abstract

Type 2 diabetes (T2D) is characterized by impaired glucose uptake in skeletal muscle and adipose tissues, which contributes to systemic hyperglycemia. GLUT4 is a crucial component in insulin-stimulated glucose uptake and its expression as well as translocation are impaired in T2D onset. This study explored the role of extracellular vesicles (EVs) derived from GLUT4-overexpressing engineered muscle constructs (G4OE-EMC) in glucose metabolism. G4OE-EMC-derived EVs enhanced glucose uptake and insulin sensitivity both in vitro, when tested on wild-type (WT) engineered muscle constructs, and in vivo using the diet-induced obesity (DIO) mouse model. Proteomic and transcriptomic analyses revealed that the EVs were enriched in IGF1 and contained reduced levels of miRNAs, such as miR-122-5p, miR-16-5p, and miR-486-5p, which target IGF1R. The multi-omic approach used here suggests a mechanism whereby G4OE-EMC-derived EVs enhance glucose metabolism via IGF1 signaling and miRNA-mediated regulation of IGF1R expression, offering a potential therapeutic strategy for T2D.

## 2. Introduction

Type 2 diabetes (T2D) and metabolic syndrome are among the most prevalent metabolic disorders, diagnosed in approximately 6% of the world population and over 10% of Americans (1). T2D is characterized by impaired glucose homeostasis and chronic hyperglycemia stemming from insulin resistance primarily in skeletal muscle, adipose and hepatic tissue (2,3).

Skeletal muscle tissue plays a pivotal role in glucose homeostasis (4). It is the site of approximately 80% of insulin-stimulated glucose uptake, rendering it crucial in maintaining blood glucose levels within a healthy physiological range. Insulin facilitates glucose uptake in skeletal muscles by promoting the translocation of glucose transporter type 4 (GLUT4) to the cell membrane via the PI3K-AKT signaling pathway, enabling glucose entry into the cell for metabolism or storage as glycogen (5). In T2D, glucose uptake in skeletal muscle tissue is markedly impaired due to insulin resistance, which is characterized by a reduced response to insulin, leading to decreased GLUT4 translocation and subsequently inadequate glucose uptake. Consequently, the patient becomes increasingly hyperglycemic, exacerbating the pathophysiological conditions associated with T2D, such as hyperinsulinemia and increased hepatic glucose production (6,7). Additionally, the muscle tissue gradually deteriorates due to glucose-starvation, driving the disease progression. The interplay between insulin resistance and the physiological state of muscle tissue is not fully understood (7,8).

The use of skeletal muscle tissue engineering as a therapeutic approach to address insulin resistance in T2D has been explored in mice. Beckerman et al. demonstrated that implantation of GLUT4-overexpressing (G4OE) skeletal muscle tissue in diet-induced obesity (DIO) mice led to overall reduction in blood glucose levels and normalization of glucose tolerance test (GTT) curves (15). Furthermore, systemic effects were observed indicated by elevated expression of myokines and proteins related to metabolic processes and reduction in liver fat levels. Additional study by our group (16) utilized a similar system in human engineered skeletal muscle tissue. We demonstrated the ability to engineer human GLUT4-overexpressing skeletal muscle with similar metabolic benefits as were shown for mouse tissue. However, the routes through which a small patch of engineered muscle can induce systemic phenomena remained largely unknown. Uncovering these routes is a crucial step towards understanding the full therapeutic potential of engineered muscle and its derived products. These results pointed towards intercellular communication between the engineered muscle and the host metabolic system.

Cell-cell communication is crucial for the regulation of metabolic homeostasis and the propagation of signals that influence insulin sensitivity and glucose metabolism (9,10). Extracellular vesicles (EVs) are key players in this process (11–13). These small, membrane-bound particles facilitate the transfer of bioactive cargo assembled from proteins, lipids and RNAs between cells to modulate cellular function in target tissues.

Micro-RNAs (miRNAs) are short non-coding endogenous RNAs that regulate gene expression by binding to their target mRNA, triggering degradation and subsequently reducing expression levels at both the transcriptomic and the proteomic levels (17). A single miRNA can regulate hundreds of targets, and a single mRNA can be targeted by several miRNAs (18). miRNAs were shown to be the most abundant RNA type in EVs (19) and have also been found to play key roles in many processes in a disease-dependent context (14,20).

This work aimed to decipher the mechanism by which engineered G4OE skeletal muscle tissue incurs systemic effects, including altered metabolic homeostasis across different systems. To this end, we isolated EVs from G4OE engineered muscle tissues, and explored their contribution to the crosstalk within the skeletal muscle tissue and other organs. We demonstrated the functional impact of the EVs on glucose metabolism in target cells both in vitro and in vivo. Proteomic profiling and small RNA analysis of G4OE-derived EVs showed enrichment of T2D-relevant signaling molecules, particularly those associated with the IGF1 signaling pathway. The results from this multi-omic analysis combined with the in vitro and in vivo evidence indicate a cell-cell communication mechanism that underlies a potential therapeutic approach for T2D.

## 2. Results

### 2.1 G4OE-EMC derived EVs display typical EV phenotypes

To examine our hypothesis that EVs play a role in the mechanism that underlies the increase in glucose uptake following G4OE-EMC implantation (Figure 1A), we characterized EMC derived EVs as indicated by the MISEV2018 guidelines (34). EVs displayed typical shape, size (Figure 1B, C) and expected protein content (Figure S1A) Specifically, proteins related to cytosolic content of EVs were 6.5 times more abundant on average in the EV fraction compared to the CM and were harvested as shown in Figure S1A (Log2 intensities_(EV)_ = 27.2, Log2 intensities_(CM)_ = 24.5, p = 0.015). EV transmembrane/anchored proteins were practically undetectable in the CM. Moreover, ribosomal protein level was 30 times lower on average in the EV proteome compared to the CM (Log2 intensities_(EV)_ = 25.5, Log2 intensities_(CM)_ = 30.4, P-value ≤ 0.001, Figure S1A). Measurement of EV sizes by dynamic light scattering (DLS) showed a reduction in size of G4OE-EMC-derived EVs compared to WT-EMC derived EVs (radius_WT_ = 122.5nm, radius_G4OE_ = 112nm, P-value < 0.01 S1B. AFM analysis corroborated these findings (radius_WT_ = 118.5nm, radius_G4OE_ = 102nm, P-value ≤ 0.01, Cohen’s d = 0.36, Figure S1C-S1D). Together, these findings indicate that the G4OE-EMC muscle construct secretes typical EVs.

**Figure 1:**
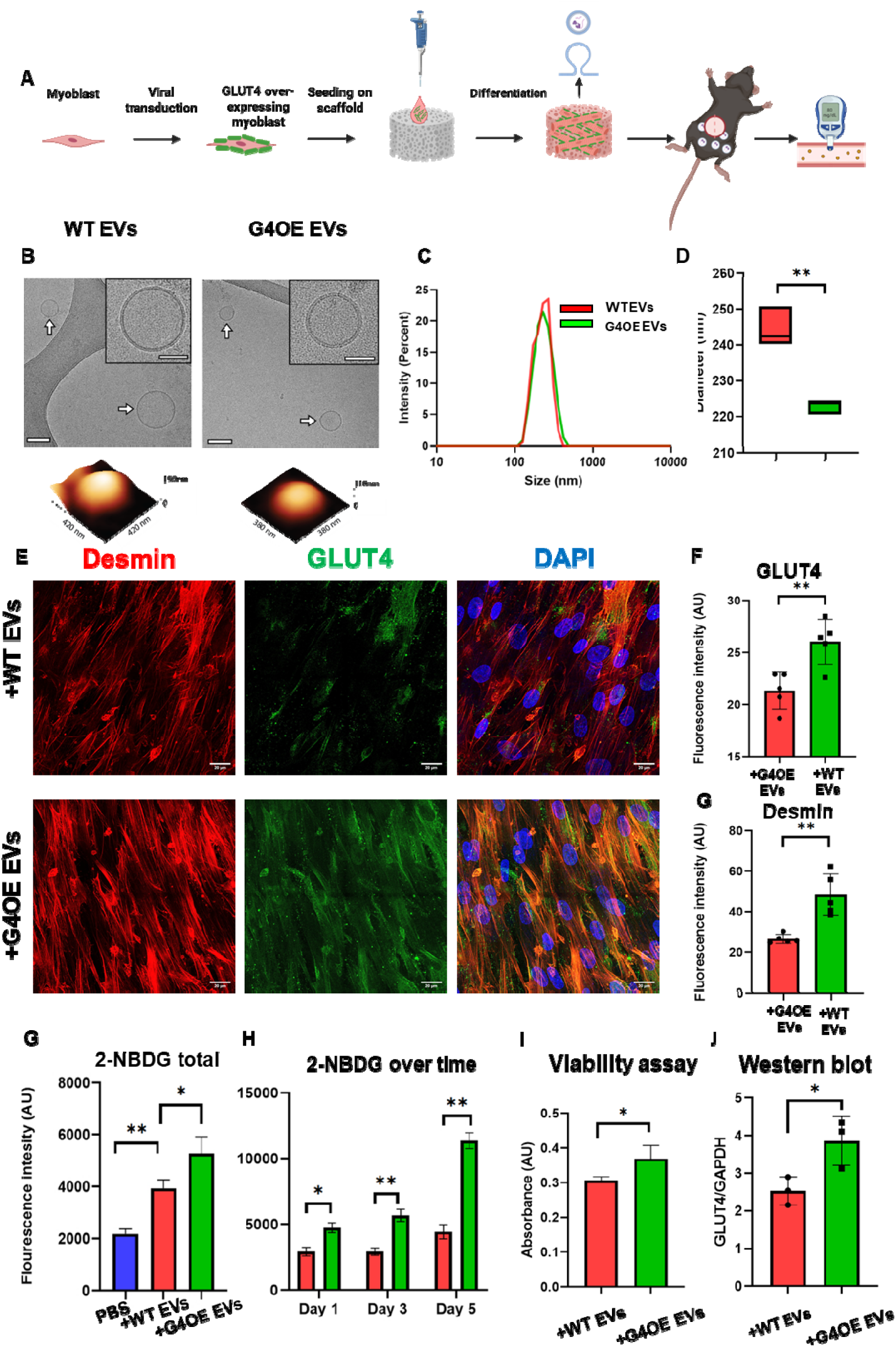
In vitro characterization of G4OE-EMC-derived EV functionality. **A)**Illustration of EMC implantation in DIO mice. B) Representative cryo-TEM analysis of WT (left) and G4OE (right) EVs (top) and their 3D shape obtained by AFM (bottom). **C)** DLS size quantification of EVs derived from WT (left) or G4OE-EMC (right). **D)** Whole-mount immunostaining for desmin (Red), GLUT4 (Green) and the nucleus (DAPI, Blue) in WT-EMC incubated with G4OE-EMC-derived EVs or WT-EMC-derived EVs. **E)** and **F)** Fluorescence quantification of GLUT4 and desmin, respectively, ****p <0.0001.**G)** 2-NBDG glucose uptake by WT-EMC incubated with G4OE-EMC-derived EVs, WT-EMC-derived EVs or PBS. Results are presented as measured fluorescence intensity. N=3. *p <0.05, **p <0.01. **H)** 2-NBDG glucose uptake by WT-EMC incubated for 1, 3 or 5 days with G4OE-EMC-derived EVs or WT-EMC-derived EVs. Results are presented as measured fluorescence intensity. N=3. *p <0.05, **p <0.01. **I)** PrestoBlue viability assay of WT-EMC incubated with G4OE-EMC-derived EVs or WT-EMC-derived EVs. N=3. *p value<0.05. **J)** Western blot quantification of GLUT4 in WT-EMC incubated with G4OE-EMC-derived EVs or WT-EMC-derived EVs. N=3, *p<0.05

**Figure 2:**
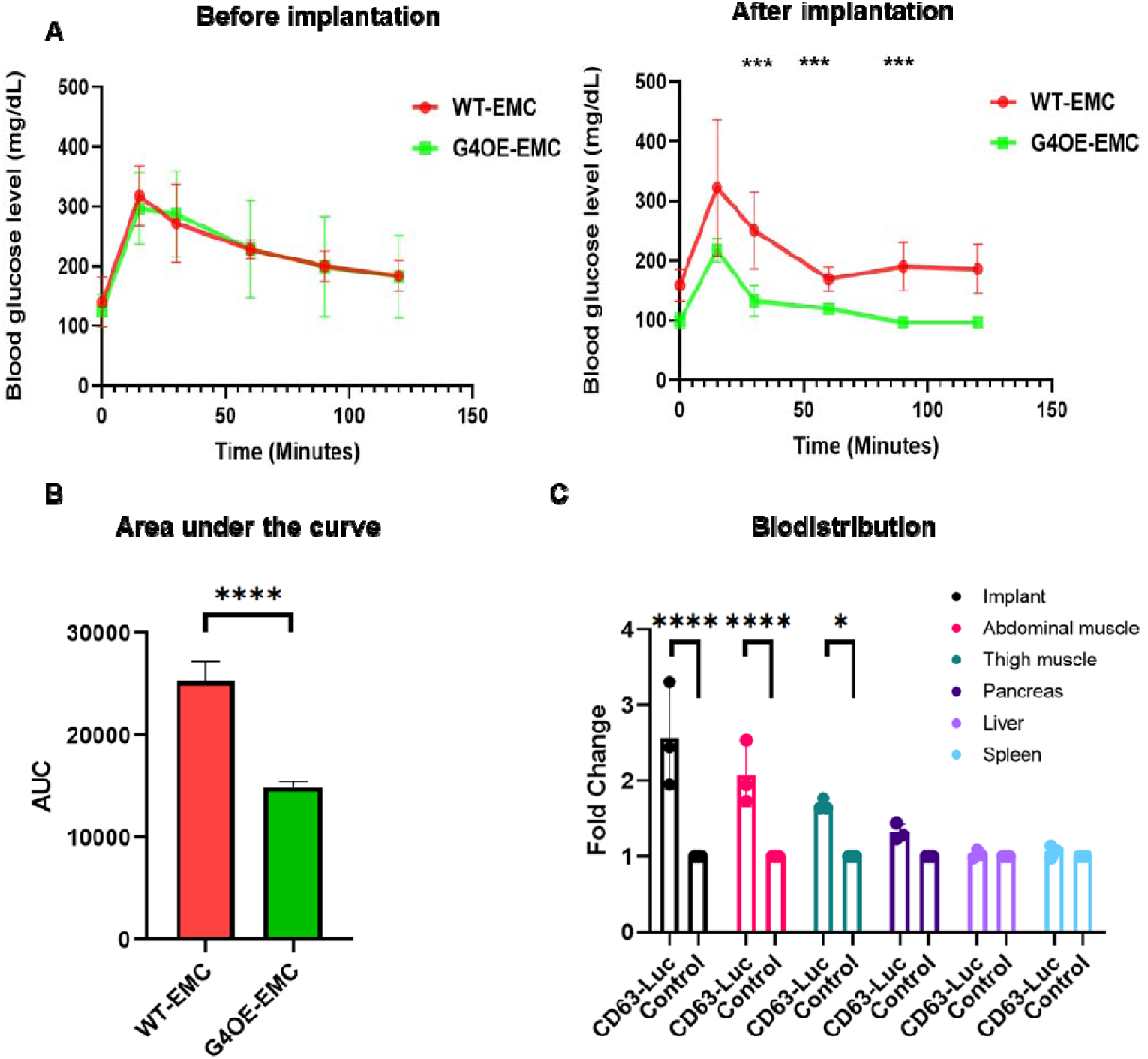
In vivo implantation of EMC. **A)** Glucose tolerance curves as measured 10 weeks after implantation of C57 G4OE-EMC (green) vs. C57 WT-EMC (red) in the abdominal muscle of DIO mice. **B)** AUC analysis of GTT curves measured after EMC implantation. N=5, *p<0.05, **p<0.01, ****p<0.0001. **C)** Optical measurements of luciferase signal distribution in different organs retrieved from CD63-luciferase-EMC-implanted mice vs. control mice implanted with an empty scaffold.

### 2.2 In vitro functionality of EMC-derived EVs

To explore the effect of G4OE-EMC-derived EVs on their target cells, a series of in vitro assays were performed to investigate glucose uptake rates and cell viability of WT-EMC in the presence of G4OE-EMC-derived EVs. WT human myoblasts were grown and differentiated on polymeric scaffolds and were administrated with G4OE-EMC-derived EVs (∼0.5*10^9 particles) for 4 days. Figure 1G shows glucose uptake of WT-EMC incubated with G4OE-EMC derived EVs, WT-EMC derived EVs and PBSX1 control. Incubation with G4OE-EMC-derived EVs resulted in ∼40% increased glucose uptake compared to constructs treated with WT-EMC-derived EVs and ∼100% increased glucose uptake compared to the PBS control. (Figure 1G). To examine the rate of increase, glucose uptake was measured every 2 days following day 1 without addition of new EVs. Our results show that the glucose uptake increased ∼2.5 fold over the course of 5 days. (Figure 1H). We conducted a PrestoBlue metabolic assay of the WT-EMCs following incubation with EVs to assess the overall metabolic state (Figure 1I). G4OE-EMC-derived EVs increased myoblast metabolism by ∼30% compared to the control (constructs incubated with WT-EMC-derived EVs). Constructs were immunostained for desmin and GLUT4 following insulin stimulation (Figure 1D-F, Figure S3). Increased GLUT4 expression and membrane localization after incubation with G4OE-EMC-derived EVs as well as increased levels of desmin, the intermediate filament of the skeletal muscle, indicated a more advanced differentiation state (Figure 1F). Western blot analysis showed a ∼50% increase in GLUT4 protein levels (Figure 1J, S2A) in myoblasts incubated with G4OE-EMC-derived EVs. Combined, the in vitro assays suggest that G4OE-EMC-derived EVs increase muscle metabolism rates, specifically GLUT4-mediated glucose uptake.

### 2.3 Implanted G4OE-EMC in DIO mice secrets EVs in vivo and causes reduction in blood glucose

To confirm EVs secretion from the engineered muscle tissue in vivo, GLUT4-overexpressing skeletal muscle cells from C57 mice strain were cultured on 3D scaffolds (GLUT4 engineered muscle constructs (G4OE-EMC)) and were implanted into the abdominal muscle of DIO mice (illustrated in Figure 1A). Glucose metabolism as indicated by fasting glucose and glucose tolerance test (GTT) measurements was assessed as previously reported (15,16). G4OE-EMC-implanted mice showed a significant decrease of approximately 40% in fasting blood glucose levels compared to WT control (Figure 3A, 3B). Moreover, they demonstrated normalization of their glucose response as shown by the GTT results, indicating partially re-established insulin sensitivity. Next, we aimed to track EVs secretion from the EMC. We produced human skeletal muscle cells (hSkMCs) engineered to secrete EVs labelled with luciferase fused to CD63. This allowed us to detect engineered tissue-derived EVs in vivo and assess their biodistribution (Figure 3C). The strongest luciferase signal (∼2.5 fold compared to the non-labelled EMC control) was found in the retrieved muscle construct. We also detected a significant signal spread to the rest of the mouse abdominal muscle (∼2 fold) and to the more distant thigh muscle (∼1.7 fold). The results indicate that EVs from the engineered construct potentially reach the entire skeletal muscle tissue of the mouse.

**Figure 3:**
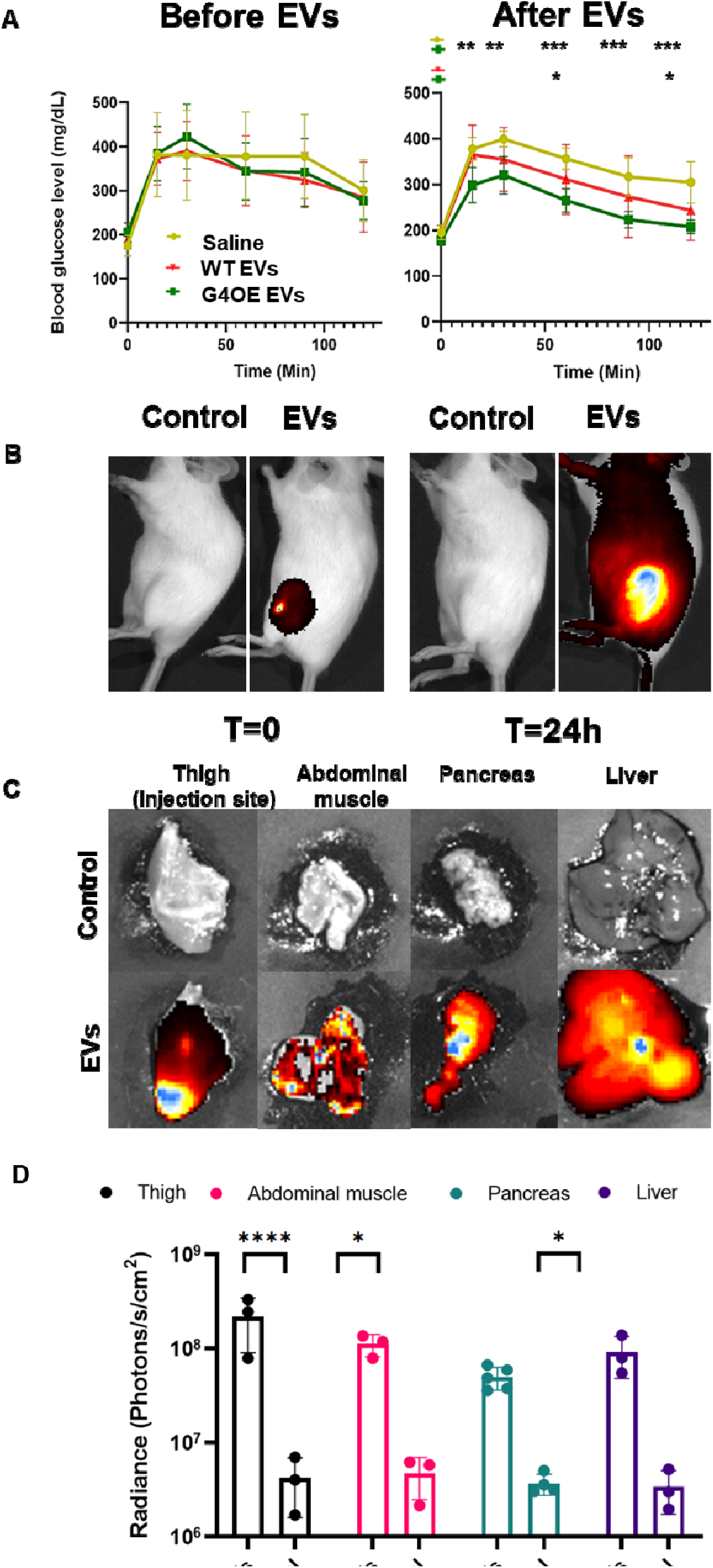
In vivo functionality and biodistribution of injected G4OE derived EVs. **A**) Glucose tolerance curves as measured for before (left) and 6 weeks after injection (right) of G4OE-EMC EVs vs. WT-EMC EVs or saline. N=7, *p<0.05, **p<0.01. ***p<0.001. **B)** Biodistribution of DiR-labelled G4OE-EMC EVs following their intramuscular injection into C57 mice thigh muscle, at the time of injection and after 24h. **C)** Representative ex-vivo images of organs extracted from animals described in (B). **D)** Fluorescence quantification of DiR-labelled EVs injected to animals described in B.

### 2.4 Injection of G4OE-EMC-derived EVs in DIO is followed by improved glucose metabolism

To characterize the impact of the G4OE-EMC-derived EVs on glucose homeostasis, WT-or G4OE-EMC-derived EVs were intramuscularly (IM) injected over the course of 10 days in 3 injections, into the thigh muscle (∼10^9 particles per injection) of DIO mice. One week after the last injection, we assessed the mice’s response to glucose challenge (GTT)for 6 weeks after injections. Figure 3A shows GTT curves prior to EVs injection and at the end point (left and right, respectively). An approximate 15% reduction in blood glucose peaks was measured in mice treated with G4OE-EMC-derived EVs compared to those treated with WT-EMC-derived EVs. Moreover, an approximate 25% reduction in glucose peaks was measured in comparison to those treated with a saline control (Figure 3A). Overall reductions in blood glucose levels as indicated by Area under the curve (AUC) are presented in Figure S2B. In addition, we aimed to assess the biodistribution of injected EVs to test our hypothesis that the systemic effect stems from intercellular crosstalk mechanisms. DiR lipophilic dye labelled EVs were injected IM to the thigh muscle and we followed the spread of the fluorescence using In Vivo Imaging System (IVIS) over 24 hours (Figure 3B,S2C). Following this, the mice were sacrificed, their organs were removed and imaged individually to detect the specific sites of EVs uptake. IVIS quantification showed that EMC-derived EVs travelled from the injection site to distant muscles (e.g., abdominal muscle) and other organs such as the pancreas and the liver (Figure 3C and 3D).

### 2.5 G4OE-EMC EVs show metabolic protein enrichment

To better understand the molecular mechanism that underlies the increased metabolism observed both in vivo and in vitro after G4OE-EMC EV treatment, the proteomic profile of the EVs was analyzed by liquid chromatography–tandem mass spectrometry (LC-MS/MS). The analysis revealed 676 proteins, of which 75% (528) were uniquely mapped to the EV proteome when compared to that of the CM (Figure 4A). A comparison between the proteome of G4OE-EMC EVs and WT EVs yielded an enrichment of 67 proteins in G4OE derived EVs (Log_2_(FC) > 0, FDR ≤ 0.05), while 22 proteins were depleted (Log_2_(FC) < 0, FDR ≤ 0.05, Figure 4B). Among these proteins, prostaglandin F2 receptor inhibitor (PTGFRN) and insulin-like growth factor 1 (IGF1) were enriched by 17-fold and 71-fold, respectively, in G4OE-EMC EVs. Gene Ontology (GO) terms enrichment analysis found that the enriched proteins were mainly related to catabolic processes, various peptidase activities and their regulation (Figure 4D). While the depleted proteins were mostly enriched for ECM-related, and RNA-binding-related proteins (Figure 4D). A similar pattern was observed when analyzing for protein-protein interactions shown in figure 5D. The proteins enriched in G4OE-EMC EVs displayed two clusters, with functions of unfolded protein binding and collagen chain trimerization found associated to metabolism, while the depleted proteins formed one cluster related to cathepsin activity. (Figure 4D). Combined, the changes in ECM organization related proteins and IGF-1, support the effect of G4OE-EMC EVs on improving muscle glucose metabolism and myogenesis.

**Figure 4:**
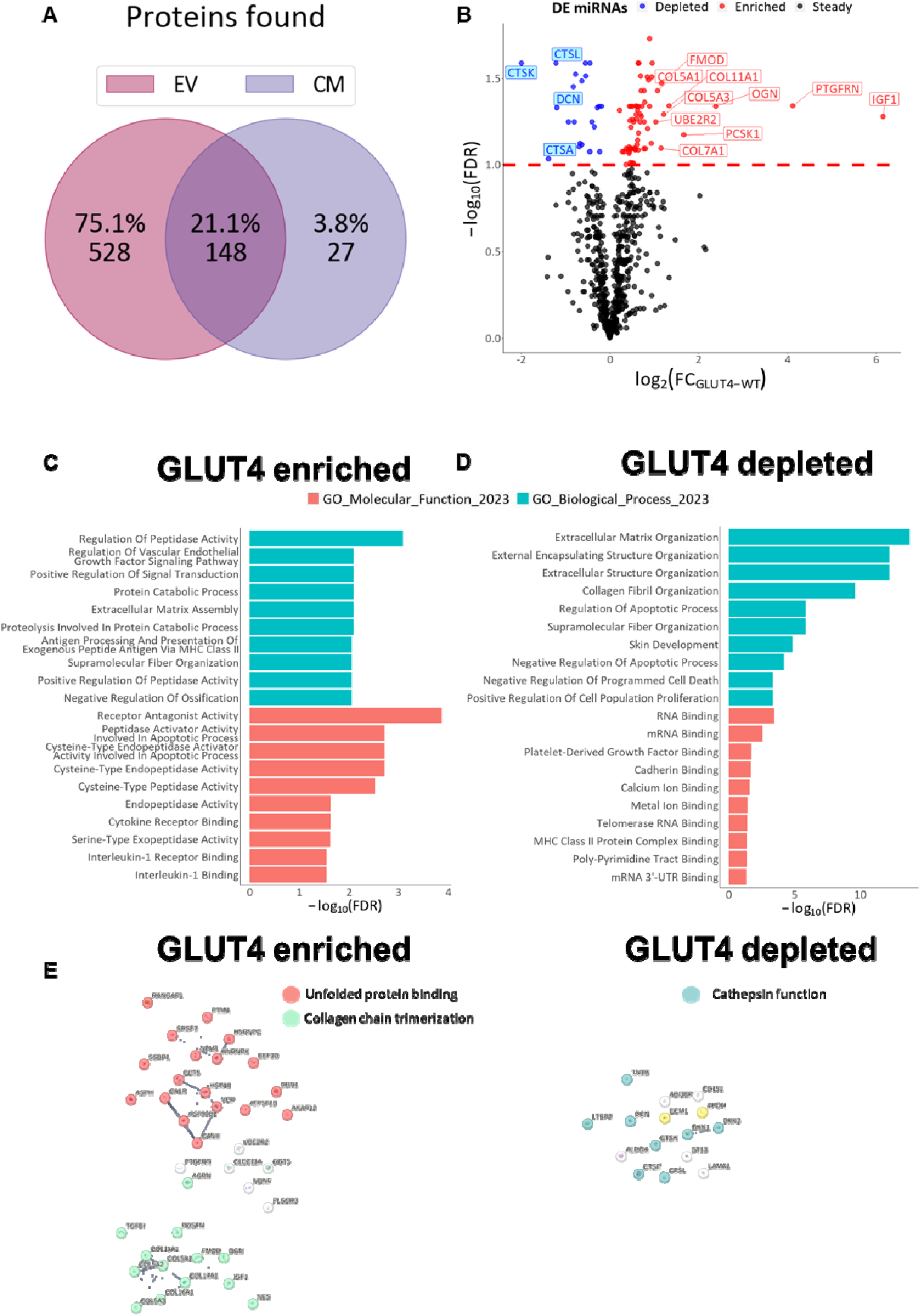
G4OE-EMC-derived EVs are enriched with metabolism-related proteins. **A)** Venn diagram of proteins found in EVs (pink) or CM (purple). **B)** Volcano plot of abundance differences of enriched (red) or depleted (blue) proteins (Log2 of the fold change, X) and their significance (-log10 of the false discovery rate, Y) in G4OE-EVs compared to WT EVs. Dashed red line denotes FDR significance threshold. **C-D)** Bar plots of top-five GO term enrichment analysis of enriched (left) or depleted (right) proteins in G4OE-derived EVs for various terms (color coded). **E)** STRING analysis for enriched (left) or depleted (right) proteins in G4OE-derived EVs. Lines denote protein interactions, thickness correlates to interaction strength and colors denote function clusters by Kmeans.

**Figure 5:**
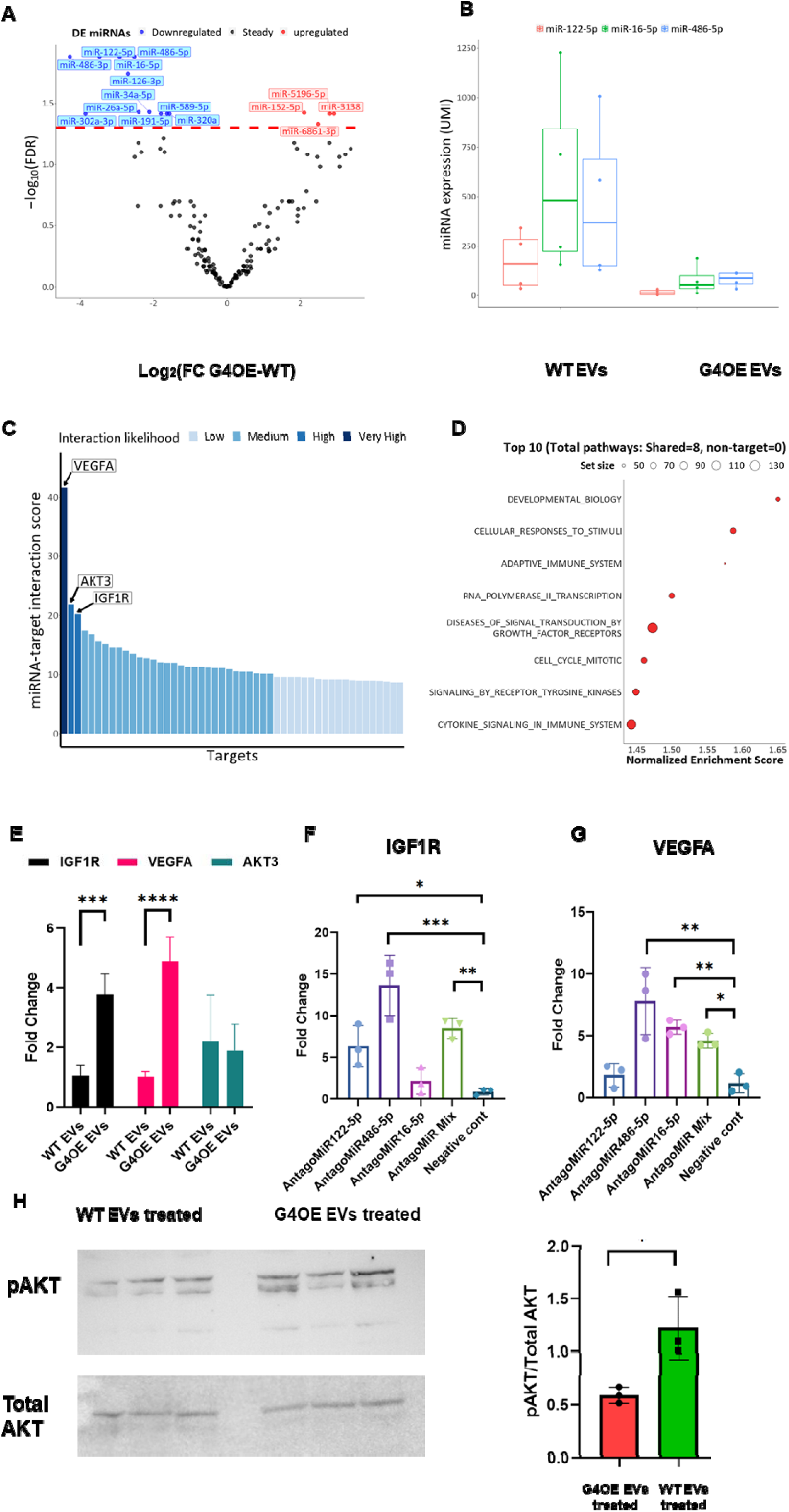
Downregulation of miRNAs targeting metabolism-related genes in the transcriptome of G4OE-EMC-derived EVs. **A)** Volcano plot for differential expression analysis of upregulated (red) and downregulated (blue) miRNAs in G4OE-compared to WT-derived EVs. EVs showing log2-fold change in expression (X) and its significance (-log10 false discovery rate, Y). Dashed red line denotes FDR significance threshold. **B)** Boxplot of expression levels in UMIs (Y) of the top most significantly changed miRNAs shown in A in G4OE and WT EVs. **C)** Bar plot of miRNA-mRNA target interaction score (Y) of miRNA shown in (B) and their targets. Category colors were assigned based on the distribution of the scores. **D)** Dot plot of KEGG gene set enrichment analysis of shared targets of the miRNAs shown in (B). X axis depicts the normalized enrichment score and gene set is denoted by point size **E)** Bar plot of differences by fold change (Y) in IGF1R (Black) or VEGFA (Pink) mRNA levels, as determined by qPCR of muscle constructs treated with G4OE or WT EVs. **F)** Fold-change differences in IGF1R mRNA levels in human myotubes following antagoMiRs transfection. **G)** Fold-change differences in VEGFA mRNA levels in human myotubes following antagoMiRs transfection. N=3, *p<0.05, ** p<0.01, *** p<0.001. **H)** Western blot of pAKT levels in muscle constructs treated with G4OE or WT EVs. N=3, *p <0.05

### 2.6 G4OE-EMC EVs regulate metabolism-related genes

The small RNA content of G4OE-EMC-derived EVs was explored by bulk RNA-seq to gain a more heuristic view of their role in regulation of muscle metabolism and development. Overall, 167 miRNAs were identified in the EVs. Among these, 15 were differentially expressed (DE, FDR ≤ 0.05) between G4OE-EMC EVs compared to WT. Only 4 (27%) DE miRNAs were upregulated, while the remaining 11 (73%) were downregulated (Figure 6A). Among these, miR-122-5p, miR-16-5p and miR-486-5p (henceforth top trio) were almost entirely depleted in G4OE-EMC EVs, with expression levels in WT EVs being 11, 7.5- and 5.8-times higher, respectively (Figure 5B). As miRNAs usually regulate multiple targets, we explored the network of genes potentially targeted by this top trio. We devised a miRNA-mRNA interaction score (IS) based on available data from the miRTargetLink 2.0 online database. Briefly, the IS considers the number of evidence for miRNA-mRNA interaction and whether these interactions are experimentally validated or computationally predicted across the three miRNAs. Higher ranking mRNAs are more likely to be targets of the three miRNAs. Interestingly, the genes most likely to be targeted by the top trio were vascular endothelial growth factor A (VEGFA), AKT serine/threonine kinase 3 (AKT3) and insulin-like growth factor 1 receptor (IGF1R) (Figure 5C). Out of 10000 bootstrap (with replacement) miRNA trios, these three genes were found to be among the top ten most probable targets in only one trio combination (p = 0.0001). Additionally, the shared target genes across the 3 miRNAs were ranked for gene set enrichment (28) according to the Kyoto Encyclopedia of Genes and Genomes (KEGG) (44) by ranking them according to their IS. The high-ranking targets were mainly enriched for signal transduction pathways and cell cycle (Figure 5D).

Given that miRNAs typically downregulate the expression of their target genes, and a reduction in the levels of these miRNAs was observed in G4OE-EMC EVs, we hypothesized that cells treated with these EVs would exhibit increased expression of the top three targets. Quantitative polymerase chain reaction (qPCR) analysis of muscle constructs treated with G4OE-EMC- or WT-derived EVs found significant upregulation of expression levels of both IGF1R and VEGFA (FC = 3.8, p-value < 0.001; FC = 4.9, p-value < 0.0001 respectively, Figure 5E). However, no change at the transcriptome level was noted for AKT. Notably, *in-vitro* silencing of miR-122-5p and miR-486-5p increased IGF1R mRNA levels in six to thirteen-fold, respectively (Figure 5F). We similarly found that MiR-486-5p and MiR-16-5p silencing increased VEGFA mRNA levels in eight and six-fold, respectively (Figure 5G). Due to its pivotal role in the cell, we assumed that changes in AKT expression levels are rare, while changes in its activation are more common (43). Western blotting for phosphorylated AKT (pAKT) identified an increase in G4OE-EMC EV-treated muscle constructs compared ^i^to those treated with WT EVs (FC = 1.6, p-value < 0.05, Figure 5H).

## 3. Discussion

Impaired glucose uptake in skeletal muscle tissue is one of the earliest detectable manifestations of T2D. EVs play a substantial role in the crosstalk required for glucose regulation, but the pathways through which they may effect this process are largely understudied. This study provided significant insights into the role of G4OE-EMC-derived EVs in modulating glucose metabolism, demonstrating their ability to enhance insulin-stimulated glucose uptake in vitro and improve glucose metabolism in vivo. Both implanted G4OE-EMCs and intramuscular injections of their EVs improved glucose tolerance and distributed systemically, reaching metabolically relevant tissues like liver and pancreas. Proteomic and miRNA analyses revealed that G4OE-EMC EVs are enriched in metabolic regulators such as IGF1, while being depleted of miRNAs like miR-122-5p and miR-486-5p that suppress its receptor IGF1R. These findings suggest a regulatory mechanism by which engineered G4OE muscle tissue can influence systemic glucose homeostasis.

One of the most heavily regulated glucose metabolism mechanisms in skeletal muscle is GLUT4 expression and translocation which drives the insulin-stimulated glucose uptake (5). This transporter is constitutively expressed and stored in designated vesicles within the cytoplasm until insulin stimulation initiates a signaling cascade resulting in GLUT4 translocation to the cell membrane and subsequent activation(4–5). Similar to Insulin, the IGF1 signaling pathway can activate similar cascades, resulting in GLUT4 translocation. The in vitro data collected in this study indicates enhanced insulin-stimulated glucose uptake and overall metabolic rate in WT engineered muscle exposed to G4OE-EMC-derived EVs.

In a previous and current studies (15,16), we showed that the effect of implanted G4OE-EMC extends far beyond locally enhanced glucose uptake, despite constituting only about 1-2% of the total mouse muscle mass. The results indicated improved insulin-mediated response to glucose challenges. Importantly, we observed similar trends following isolated EVs injection into DIO mice. Past studies reported that CM of insulin-resistant muscle tissue can induce diabetic phenotypes in healthy muscle cells (46). In this context, the role of EVs in driving the diabetic pathological mechanisms has also been investigated (15). Biodistribution analysis of both EVs secreted by the implanted tissue and of isolated EVs injected into the mice’s thigh muscle indicates that EVs travel across the skeletal muscle tissue, reaching distant muscles in the host. This communication within the skeletal muscle system plays a crucial role in maintaining glucose homeostasis (3–5). Interestingly, we observed that WT-EMC derived EVs were consistently about 40 nm larger in diameter compared to G4OE-EMC derived EVs. Evidence in the literature suggests easier and more efficient uptake of smaller EVs, thereby enhancing their potential biochemical and therapeutic effect (8).

Our multi-omics analysis implied that overexpression of GLUT4 led to an increase in metabolic pathways relevant for muscle development and functionality. G4OE-EMC EVs featured IGF1-enriched content, which, in turn, increased AKT activation via phosphorylation in host cells. These results are in accordance with past studies showing that IGF1 is an upstream activator of AKT (50). Recently, Dreher et al (55) found that supplementation of IGF1 to the culture medium improved in vitro differentiation, glucose metabolism and response to electrical stimulation. These results support prior evidence of the effect of IGF1 on muscle differentiation and subsequent metabolic functionality (56,57). Notably, the proteomic analysis also indicates the WT derived EVs contain significantly higher levels of Cathepsins compared to the G4OE derived EVs. In recent years it was found that Cathepsins are elevated in T2D patients (7,69). In light of this, G4OE-EMC EVs may impact neighboring muscle by activating the IGF-1 signaling pathway resulting in increased GLUT4 expression and translocation. This is further supported by the increase of IGF1R mRNAs observed in G4OE-EMC-derived EVs-treated recipient muscle constructs. The effect may be attributable to reduced levels of miR-122-5p, miR-16-5p and miR-486-5p miRNAs in the G4OE-EMC derived EVs to almost no expression compared to WT-EMC derived EVs as these miRNAs were previously shown to regulate IGF1R in hepatic cells ().

In addition, VEGFA expression was increased in G4OE-EMC EVs compared to WT EVs. Like IGF1, VEGFA is an activator of AKT (54) and may contribute through a parallel pathway to GLUT4 translocation to the membrane, resulting in increased metabolism and glucose uptake. The analysis results were further validated by inhibiting the three MiRs using antagoMiRs in WT human skeletal muscle cells and examining the inhibition effect on IGF1R and VEGFA expression levels.

Intriguingly, the three downregulated miRNAs identified in the current work were shown to play a role in muscle differentiation and function (59–61). In particular, MiR-486-5p has been shown to regulate IGF1R and other components of the IGF1 signaling pathway as well as VEGFA in the context of skeletal muscle system (62–64). Furthermore, as muscle tissue is hypoxic, its function and differentiation require high amounts of oxygen and nutrients, which are supplied through increased vascularization (5). Indeed, mRNA of VEGFA, a well-documented vascularization factor (6), was upregulated in G4OE-EMC EVs and may indicate increased muscle function or differentiation.

Despite the aligning multi-omics results, it must be acknowledged that there may be other pathways affecting glucose metabolism. While this work focused on EVs secreted from engineered tissue, many factors, particularly myokines, are secreted directly rather than through vesicles. For instance, as demonstrated in our previous work, the myokines IL-8 and IL-10 may play a role in the systemic metabolic effect produced by the engineered tissue (15). Furthermore, while the top three mRNA targets were most likely targeted by one trio combination out of 10000, other combinations of down regulated miRNAs may result in unique targets of their own. On the other hand, less significantly differentiated miRNAs are more likely to be the result of noise in the experimental system and are less likely to yield relevant results. Future studies will be needed to evaluate other potential miRNA candidates and their subsequent effect based on our suggested approach. Moreover, different isolation methods yield different EV populations (24,33) and their possible effect in this model ought to be studied.

Our findings suggest a possible mechanism of glucose regulation. The G4OE-EMC-derived EVs are likely to act through IGF1 signaling and possibley miRNA modulation, specifically by downregulating miRNAs that target IGF1R, thereby boosting IGF1 receptor expression and facilitating enhanced glucose uptake. This intercellular communication pathway emphasizes the systemic potential of G4OE-EMC as a therapeutic strategy to address insulin resistance.

## 4. Methods

### 4.1 Cell culture

Human skeletal muscle cells (hSkMC) were purchased from ScienCell (cat #3520). The cells were maintained in growth medium for the proliferation stage (ScienCell, Cat #3501) and then transferred into differentiation medium (high-glucose DMEM containing 1% penicillin and streptomycin (P/S), 1% Glutamax and 3% horse donor serum).

### 4.2 GLUT4 overexpression

To generate GLUT4-overexpressing (G4OE) hSkMCs, T293 cells (ATCC) that were maintained in full DMEM were first transduced with the pGenlenti lentiviral plasmid containing the GLUT4 cDNA and lentiviral particles were produced using the Lenti-X universal packaging system (Takara Bio, USA). hSkMCs were seeded at a concentration of 4x10^5^ cells/well of a 6-well plate and incubated the following day with viral particles in the presence of polybrene (6 μg/ml) (Sigma-Aldrich), for 24 h. The transfection medium was then replaced with fresh medium and the cells were incubated for 48 h to allow expansion. Stably expressing cells were positively selected using puromycin antibiotic (1.5 μg/ml).

### 4.3 CD63-luciferase cell transduction

To generate CD63-luciferase-expressing hSkMCs, T293 cells were first transduced with the pGenlenti lentiviral plasmid containing the CD63 fused to ThermoLuc sequence cDNA, and lentiviral particles were produced using the Lenti-X universal packaging system (Takara Bio, USA # 631278). hSkMCs were seeded at a concentration of 4x10^5^ cells/well of a 6-well plate and incubated the following day with viral particles in the presence of polybrene (6 μg/ml) (Sigma-Aldrich TR-1003), for 24 h. The transfection medium was then replaced, and the cells were incubated for 48 h to allow expansion. Stably expressing cells were positively selected using puromycin antibiotic (1.5 μg/ml) (Sigma Aldrich P7255). The luciferase signal was detected by injection of 15 mg/ml D-Luciferin substrate (BIOSYNTH) prepared in saline.

### 4.4 Scaffold synthesis

Salt leaching technique was used to create porous biodegradable scaffolds out of PLLA (Polysciences, Warrington) and PLGA (Boehringer Ingelheim). The constructs have a pore size of 212 to 600 μm and 93% porosity as was previously reported (15). From each polymer, a 5% (w/v) solution was prepared separately by dissolving 0.5 g of polymer in 10 ml of chloroform (Bio-Lab Ltd). The polymers were mixed in a 1:1 ratio to create a PLLA/PLGA solution. NaCl (0.4 g) was dissolved in 0.24 ml of PLLA/PLGA solution in Teflon cylinder molds (18 mm internal diameter). The Teflon molds were left overnight to allow chloroform evaporation. The following day, scaffolds were removed from the molds and placed into histology cassettes. The salt was leached by washing with distilled water on a magnetic stirring plate; the water was exchanged every hour for 6 to 8 hours. The scaffolds were dried and frozen overnight at −80°C. The following day, the scaffolds were lyophilized 24h and kept dry under vacuum until use. One day before seeding, round pieces of scaffold material of 6-mm diameter were prepared using a biopsy punch (Miltex) and sterilized in 70% ethanol overnight. Prior to seeding, the scaffolds were washed twice in PBS 1× (Sigma-Aldrich) and dried using vacuum.

### 4.5 Human skeletal muscle tissue engineering

hSkMC were cultured in a T-150 flask (FroggaBio) to 70% confluency in proliferation medium, washed with PBS, trypsinized with 2X trypsin-EDTA (Biological Industries), aliquoted according to the number of scaffolds, centrifuged at room temperature (RT) (1200 rpm, 4 min) and then resuspended in 10 μl of a 1:1 mixture of fibrinogen (15 mg/ml) (Sigma-Aldrich) and thrombin (20 U/ml) (Sigma-Aldrich). The cell suspension, 0.75 × 10^6^ cells per scaffold, was seeded onto PLLA/PLGA scaffolds and incubated (40 min, 37 °C, 5% CO_2_). Then, 2 ml hSkMC growth medium with 10% fetal bovine serum (FBS) and 1% P/S, were added to each well. After 2 days of incubation, the scaffolds were transferred to a new plate using sterile forceps and 2 ml differentiation medium (DMEM, 5% horse serum, 1% P/S and 1% Glutamax) were added to each well. Constructs were cultured in differentiation medium for three weeks.

### 4.6 Myogenic EV isolation

Myogenic EVs were isolated from the conditioned medium of engineered skeletal muscle constructs. To this end, engineered muscle constructs were transferred to EV-depleted differentiation medium produced using the FBS exosome depletion kit (Norgen Biotek). Conditioned medium was collected 48 h thereafter and filtered through a 0.45 μm syringe filter and centrifuged for 10 min, 2000 x g. EVs were isolated using the exoEasy Maxi Kit (Qiagen) according to the manufacturer’s instructions.

### 4.7 Dynamic light scattering (DLS)

Size and concentration of isolated EVs were measured using a Malvern Zetasizer ultra (Malvern, UK).

### 4.8 Transmission electron cryomicroscopy (Cryo-TEM)

3 μl of EV sample solution was applied to the carbon side of Lacey Carbon 300 mesh copper grid (Ted Pella, USA), glow discharged for 120s at 0.5 mbar and 18 W with Evactron CombiClean System (XEI Scientific, USA). Then the grids were blotted for 1.5 s from both sides at 4°C and 100% humidity and plunge frozen into precooled liquid ethane using Vitrobot Mark IV (Thermo Fisher Scientific, USA). Cryo-TEM images were collected using Talos Arctica TEM/STEM, equipped with Falcon 4i direct detector (Thermo Fisher Scientific, USA) and operated at 200 kV. Image acquisition was performed using Velox 3.14 software (Thermo Fisher Scientific, USA) at a nominal magnification of 45.000× (pixel size 0.62 nm, underfocus 6-10 µm) or 120.000× (pixel size 0.118 nm, underfocus 3-5 µm) with a total dose not exceeding 100 e-/Å^2^

### 4.9 Atomic force microscopy (AFM)

Freshly cleaved mica surface was incubated with 10 mM MgCl_2_ solution for 2 min and then rinsed with 200 μl PBS. EVs solution (50 μl) was placed on the Mg-modified mica for 10 min, after which 3 ml PBS was added before scanning with a JPK Nanowizard III AFM microscope (Berlin, Germany) in QI mode using qp-BioAC-Cl-CB-3 probe, spring constant≈0.06 N/m (Nanosensors). Image analysis was performed using Gwyddion or JPK-SPM data processing software. Particle size analysis was conducted using grain analysis in Gwyddion (20). The size of each particle was measured as the equivalent disc radius, (the radius of the disc with the same projected area of each particle).

### 4.10 In vitro glucose uptake

To determine basal (i.e., GLUT1-mediated) or insulin-stimulated (i.e., GLUT4-mediated) glucose uptake, differentiated myotube cells in either 24-well or 96-well plates were transferred to low-glucose differentiation buffer consisting of low-glucose DMEM, 1% P/S and 2% donor horse serum, ∼16 h before the experiment. Cells were then washed with warm D-PBS+ and incubated (37 °C, 5% CO_2_, 3 h) in Krebs-Ringer (KRPH) buffer. The buffer was replaced by KRPH buffer supplemented with 200 μM of the fluorescent, non-metabolizable glucose analogue 2-(N-(7-nitrobenz-2-oxa-1,3-diazol-4-yl)amino)-2-deoxyglucose (2-NBDG), and 100 nM insulin (30 min, 37 °C, 5% CO_2_). The reaction was terminated by washing the cells three times with cold PBSX1. Constructs were lysed with RIPA buffer (Sigma-Aldrich) supplemented with 1% protease inhibitors cocktail (Sigma), homogenized with a IKA™ T 10 Basic ULTRA-TURRAX™ homogenizer and centrifuged for 5 min at maximum speed. The supernatant was collected and fluorescence was measured with a plate-reader fluorometer at an excitation wavelength of 465 nm and emission wavelength of 540 nm.

### 4.11 PrestoBlue metabolism assay

PrestoBlue (Thermo Fisher A13261) assay was performed according to the manufacturer’s instructions. Briefly, the reagent was added to the muscle constructs culture medium in a 1:10 ratio. Then, the constructs were incubated for 90 min at 37°C in a 24-well plate under shaking, after which the supernatant was transferred in triplicates to a new 96 well plate. Absorbance was recorded at 562 nm using 595 nm as the reference wavelength. Percent of reduced PrestoBlue was calculated according to the formula provided by the manufacturer.

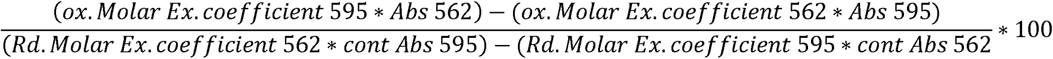

### 4.12 Whole-mount immunostaining

hSkMC-seeded scaffolds were cultured for 1-3 weeks, washed with PBS, fixed in 4% paraformaldehyde for 10 min, and washed again with PBS. Next, the scaffolds were permeabilized with 0.3% Triton X-100 for 10 min, washed with PBS and soaked in a blocking solution (5% BSA in PBS, 2 h, RT). Scaffolds were then incubated with a primary antibody diluted in blocking solution (overnight, 4 °C). The following day scaffolds were rinsed with PBS and incubated with secondary antibodies diluted in PBS mixed with 4’,6-diamidino-2-phenylindole (DAPI) (1:1000; Sigma) (2 h, RT). Samples were then washed with PBS and stored at 4CC until imaging. Proteins were stained using the following primary antibodies: rabbit anti-desmin (1:200 Abcam), mouse anti-MYH (1:50 R&D systems), mouse anti-GLUT4 (1:100 Invitrogen). Secondary antibodies: Alexa fluor 647 donkey anti-mouse (1:400; Jackson, USA), Alexa fluor 488 donkey anti-rabbit (1:400 Jackson).

### 4.13 Western blot

Scaffolds (triplicates) were washed with PBS and homogenised in RIPA solution (Sigma-Aldrich) supplemented with a protease inhibitors cocktail (100 mM PMSF, 2 mg/ml leupeptin, 50 mM Na-orthavanadate, 50 mM aprotinin (Sigma-Aldrich)). Total protein content was determined using the Bradford method (Bio-Rad, Hercules, CA). Proteins were separated on a 10% SDS-PAGE gel (NuPage©, Life Technologies) and transferred to a nitrocellulose membrane (Bio-Rad, Hercules, CA). After blocking with 5% non-fat milk (Bio-Rad, Hercules, CA), the membranes were immunoblotted (overnight, 4 °C) with mouse anti-GLUT4 (Invitrogen), rabbit anti-tAKT, rabbit anti-pAKT ser473 and mouse anti-GAPDH (Santa Cruz) antibodies, diluted 1:1000 in blocking buffer. The bound antibodies were detected following incubation with horseradish peroxidase-conjugated anti-mouse IgG or anti-rabbit IgG diluted 1:5000 (GE Healthcare Life Sciences), followed by incubation with the enhanced chemiluminescence Western blotting reagent (ECL; Amersham Biosciences). Images were acquired with the LAS-3000 imaging system (FujiFilm).

### 4.14 Quantitative polymerase chain reaction

Quantitative polymerase chain reaction (qPCR) was performed using the TaqMan Fast Advanced Master Mix (Applied Biosystems, Thermo Fisher Scientific) according to the manufacturer’s instructions. The TaqMan Gene Expression Assay (Thermo Fisher Scientific) primers for IGF1 receptor (IGF1R), vascular endothelial growth factor A (VEGFA) and glyceraldehyde-3-phosphate dehydrogenase (GAPDH) were used. The reaction was performed in the QuantStudio1 (Applied Biosystems, Thermo Fisher Scientific). The results are presented as 2^−ΔΔct^ and normalized to the GAPDH housekeeping gene.

### 4.15 Diet-induced obesity model

All animal procedures were performed according to protocols approved by the Institutional Animal Care and Use Committee of the Technion Israel Institute of Technology. Male C57BL6 mice, aged 7-8 weeks, were monitored for 2 weeks for basal weight and fasting glucose levels were measured twice a week. Food was withheld from mice for 6 h and glucose levels were determined using a glucometer (Freedom Lite, FreeStyle). After establishing basal levels, the mice were transferred from regular chow to high-fat diet (HFD) chow (Teklad) with 60% fat content. Fasting glucose blood levels and weight were monitored once a week during the 12 weeks of diabetes induction.

### 4.16 Engineered tissue implantation

The engineered constructs were implanted after an in vitro culturing period of 2-3 weeks. Mice were anesthetized with a ketamine (100 mg/kg body weight; Vetoquinol) and xylazine (10 mg/kg body weight; EuroVet), intraperitoneally delivered using a 27 G needle, or with 2.5% isoflurane (Abbott). A portion of fur around the animal’s abdomen was removed by shaving and depilation cream. A small incision was made allowing access to the linea-alba and surrounding tissue, where a 2- to 3-mm full-thickness tissue section was removed from the abdominal muscle. The constructs were sutured into the defect margins with 8-0 silk sutures (Assut Sutures). The outer skin was closed and sutured using 5-0 AssuCryl surgical sutures (Assut Sutures). Mice were monitored closely for 2 h to ensure full recovery.

### 4.17 EVs IM injection

Purified EVs were diluted to 5*10^9 particles/ml in saline and 50 µll were injected into the thigh muscle using a 27G syringe needle. In total, three injections were administered at 3-day intervals.

### 4.18 Glucose tolerance test

A GTT was performed weekly, starting from 7 days and up to 2 months following EV injections. After a 6-hour fast, a 10% w/v glucose solution (B Braun) was intraperitoneally injected into both control and diabetic animals. Blood glucose was measured with a glucometer (FreeStyle, Abbot) 0, 15, 30, 45, 60, 90 and 120 min following glucose administration.

### 4.19 Biodistribution

Injected EVs were labelled by incubation with 2.5 µM DiR (Invitrogen) in PBS for 1 h at 37 CC under shaking, followed by 3 washes with PBS filtered through 0.02 µm filter, using a 100 kDa cutoff centricon (millipore). The EVs were subsequently injected into the mice’s thigh muscle using a 27 G syringe needle. The mice were imaged using a PerkinElmer IVIS Spectrum CT at T=0, 1 h, 24 h. Following the final time point, the mice were sacrificed and organs were removed for imaging.

### 4.20 Small RNA next-generation sequencing

Total RNA was extracted from EVs that were suspended in ExoEasy elution buffer using the miRNeasy Micro Kit as per the manufacturer’s protocol (Qiagen, #217084), with optimization for EV RN isolation. The modifications included use of Trizol-LS (instead of Qiazol) with an EV elution buffer: Trizol-LS at 1:3 ratio, following the addition of chloroform at 0.8 volume of EV elution buffer. RNA was quantified with a Qubit fluorometer and the RNA Broad Range Assay Kit (Thermo Fisher Scientific, #Q10211). For small RNA next-generation sequencing, libraries were prepared from 7.5Cng total RNA, when the concentration was measurable, and otherwise, from 5 µll RNA, using the QIAseq miRNA Library Kit, according to the manufacturer’s protocol (Qiagen, #331502). Samples were randomly allocated to library preparation and sequencing in batches of up to 48 samples based on QIAseq miRNA NGS 96 Index IL UDI-F (Qiagen, #331945) by an experimenter who was blinded to the identity of samples. Library concentration was determined with a Qubit fluorometer (dsDNA High Sensitivity Assay Kit, Thermo Fisher Scientific, #Q32854) and library size with TapeStation D1000 (Agilent). Libraries with different indices were multiplexed and sequenced on a NextSeq 500/550 v2 (Illumina, #20024906) or a NovaSeq 6000 flow cell (Illumina, #20028401), with 75-bp single read or paired end reads with 6-bp index. FASTQ files were de-multiplexed using the user-friendly transcriptome analysis pipeline (UTAP) (21). Human miRNAs and variants, as defined by miRBase - V22 (22,23), were mapped using Qiagen’s RNA-seq analysis portal (24). miRNA variants were considered as up to 2 bp changes (insertion, deletion, shift and mismatch) up/downstream from MirBase canonical sequence, with a maximum of 2 mismatches allowed in alignment.

### 4.21 EV-derived miRNA differential expression analysis

Mapped reads were filtered based on average unique molecular identifier (UMI) expression of at least 25 across technical repeats in at least one condition. Remaining miRNAs were subjected to differential expression analysis, performed with the DEseq2 package (24). Cook’s distance (25) and principal component analysis were run to determine technical repeat outliers. p values of differential expression analysis were corrected to FDR (26) and a threshold of 0.1 was set to define significance.

### 4.22 miRNA target analysis

Selected miRNAs were queried for their mRNA targets in the miRTargetLink 2.0 online database (27). For each mRNA, targets were ranked by the following interaction score (IS): targets were assigned to one of three categories based on miRTargetLink annotation - predicated (no experimental validation), validated and strongly validated. Number of evidence for each mRNA-miRNA interaction were counted across miRNAs in each category. The sum of predicted targets was penalised by dividing it with the mean count (1.81) of validated targets across all miRNAs. Strongly validated targets were rewarded with the addition of their count multiplied by mean count of validated targets. If mRNA target was validated but not strongly, no score was added. Finally, scores were summed for each mRNA target across categories. Resulting ranked genes were subjected to gene set enrichment analysis (GSEA) by “fgsea” package in R^28^ and gene sets downloaded from GSEA MSig-DB website (29,30). Targets were assigned to categories based on the distribution of the target scores.

### 4.23 Proteomic sample preparation

EV samples were suspended in the ExoEasy Maxi Kit extraction buffers, which were adjusted to final concentrations of 5% SDS, 100 mM Tris-HCl (pH 8.0), and 10 mM DTT. The samples were heated at 95 °C for 5 minutes and sonicated twice to ensure complete protein denaturation and solubilization. Proteins were then precipitated with acetone, and the resulting pellets were resuspended in a solution containing 8 M urea, 400 mM ammonium bicarbonate, and 10 mM DTT. For protein reduction, the samples were incubated at 60 °C for 30 minutes. Alkylation was performed by adding 35.2 mM iodoacetamide in 100 mM ammonium bicarbonate and incubating at room temperature for 30 minutes in the dark. The urea concentration was diluted to 1.5 M using 66 mM ammonium bicarbonate before adding modified trypsin (Promega) at an enzyme-to-substrate ratio of 1:50 (w/w). Digestion was carried out overnight at 37 °C. A second digestion step was performed by adding more trypsin at a 1:100 (w/w) enzyme-to-substrate ratio and incubating for an additional 4 hours at 37 °C to ensure complete proteolysis. Tryptic peptides were desalted using homemade C18 StageTips, dried under vacuum, and reconstituted in 0.1% formic acid. Peptide separation was conducted using an Easy-nLC 1200 system (Thermo Fisher Scientific) coupled to a homemade capillary column (30 cm length, 75 μm inner diameter) packed with Reprosil C18-Aqua resin (Dr. Maisch GmbH, Germany). The peptides were loaded in solvent A (0.1% formic acid in water) and eluted with a linear gradient from 6% to 34% solvent B (80% acetonitrile with 0.1% formic acid) over 60 or 180 minutes, followed by a gradient from 34% to 95% solvent B over 15 minutes, and maintained at 95% solvent B for an additional 15 minutes at a flow rate of 0.15 μL/min.

Mass spectrometric analysis was performed on an Exploris 480 mass spectrometer (Thermo Fisher Scientific) operating in positive ion mode. Full MS scans were acquired over an m/z range of 300–1500 or 350–1200 with a resolution of 60,000 for MS1 and 15,000 for MS2. Data-dependent acquisition (DDA) was employed to select the top 20 or 30 most intense ions with charges greater than one for fragmentation. High-energy collision-induced dissociation (HCD) was used with a normalized collision energy of 27%. Automatic gain control (AGC) targets were set at 3×10^6 for full MS scans and 1×10^5 for MS/MS scans. The intensity threshold for triggering MS/MS events was 1×10^4. A dynamic exclusion of 20 seconds was applied to minimize repeated sequencing of the same ions.

### 4.24 Proteomic analysis

MaxQuant software version 2.1.1.0 (1) was used for peak-picking from the mass spectrometry data and the Andromeda search engine was used for protein identification, while searching against the human proteome UP000005640 from the Uniprot database (form Jan 2022, 81,803 entries), with mass tolerance of 6 ppm for the precursor masses and 20 ppm for the fragment ions. Oxidation on methionine and protein N-terminus acetylation were accepted as variable modifications and carbamidomethyl on cysteine was accepted as a static modification. Minimal peptide length was set to seven amino acids and a maximum of two miscleavages was allowed. The data were quantified by label-free analysis using the same software. Peptide- and protein-level FDRs were filtered to 1% using the target-decoy strategy. The protein table was filtered to eliminate the identifications from the reverse database and common contaminants, and single peptide identifications. Statistical analysis of the identification and quantization results was done using Perseus 1.6.15.0 software (31)

### 4.25 EV-derived proteomic profiling

Intensity values were log_2_-transformed and proteins with missing values were removed. The quality of EVs extracted from the control line was validated according to the MISEV2018 guidelines (32) and compared to the profile of the conditioned media (CM). Consecutively, remaining missing values were imputed by estimating the mean and standard deviation of each protein across all repeats per condition and randomly sample a value from a normal distribution (random seed set to 42) with the protein’s standard deviation and a mean smaller by 1.75 standard deviations. Mean differences of protein levels between groups were compared across proteins with an FDR threshold of 0.1 deeming significance. Significantly enriched/depleated proteins were submitted to enrichment analysis by Enrichr (33). Enrichment p-values were corrected by FDR and a threshold of 0.05 was set for significance. Protein-protein interactions (PPI) were calculated by the STRING V12.0 (34) software and were clustered by Kmeans (k = 2).

### 4.26 Statistical analysis

Data were analyzed using R-V4.0.4 (35), Python-V3.9.2 (36) and GraphPad software. miRNA analyses were run in R-V4.0.4 (35). Differential expression analysis was conducted using the “DEseq2” package (24) and corrected for multiple hypotheses by FDR. To determine the significance of shared miRNA targets, Monte-Carlo simulations (37) were run on ∼30k random trio combinations and the interaction score per target was calculated and sorted from highest to lowest. The occurrence of each of the top-three original targets was counted among the top 10 targets of each trio. The number of trios in which occurrences of all top-three targets appeared was divided by the number of permutations to form a p value. Gene set enrichment analysis was conducted using the “fgsea” package (28) with minimum/maximum set size of 15/500, respectively. Gene sets were collapsed together based on similarity using the collapsePathways function. Proteomic analyses were run with Python-V3.9.2(36). Differences between protein levels and between EV sizes were compared by student t-test (38) assuming equal variance. Cohen’s d (39) was used to measure effect sizes. For protein level comparisons, p-values were corrected for FDR when comparing G4OE derived EVs to WT derived EVs. Enrichment analysis was conducted using the “gseapy” Python package (40). ANOVA (41) was used to test differences in MISEV2018-related proteins between the EV fraction and CM following Tukey HSD post-hoc analysis (32,42). miRNA and protein figures were plotted using R.

The applied statistical tests used to determine group differences were unpaired Student’s *t* test for qPCR assays, staining quantification and PrestoBlue analysis, an ordinary one-way analysis of variance (ANOVA) for the 2-NBDG assay, and area under the curve (AUC) for the GTT assay. Two-way ANOVA was performed for GTT assays and normalization of glycemic value follow-up in DIO mice. p values below 0.05 were taken to indicate a statistically significant difference. Data are presented as means ± SD.

## Acknowledgments

We would like to thank D. Safina for editorial assistance with the preparation of this manuscript We also thank the Technion’s Preclinical Research Authority and M. Tendler for professional animal care; and the Technion’s “Smoler Proteomics Center” for support with proteome measurements and analysis.

## Footnotes

i F,/ fvd,/ .0072.

