## Supplementary Information for "Extracellular Vesicles Mediate Glucose Regulation by GLUT4-Overexpressing Engineered Muscle Tissue in T2D Mice"

**Supplementary Figures:**

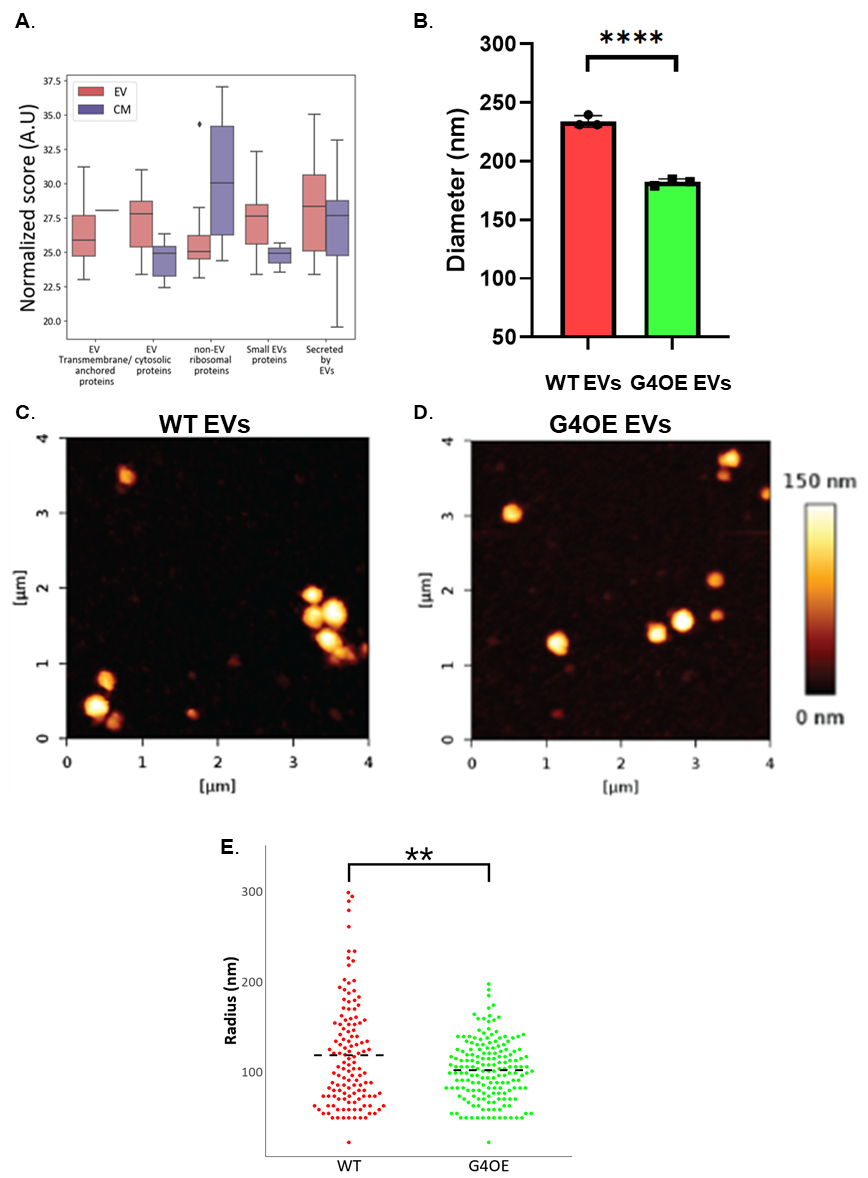

**Figure S1:** **Size comparison of EVs by AFM and DLS.**

**A)** Boxplot of levels of MISEV2018 guidelines protein groups derived from G4OE-EMC EVs (purple)/CM (pink). Y axis denotes log_2_ of protein level intensities. **B)** DLS distribution of diameters of EVs derived from G4OE (224 nm) and WT muscles (245 nm). **C and D)** Representative AFM analysis 10 µm resolution images of WT EVs (C) and G4OE EVs (D). **E)** Besswarm radius in nm (Y) of EVs derived from WT and G4OE muscles (X), as quantified by AFM. Dashed line denotes the average EV radius.

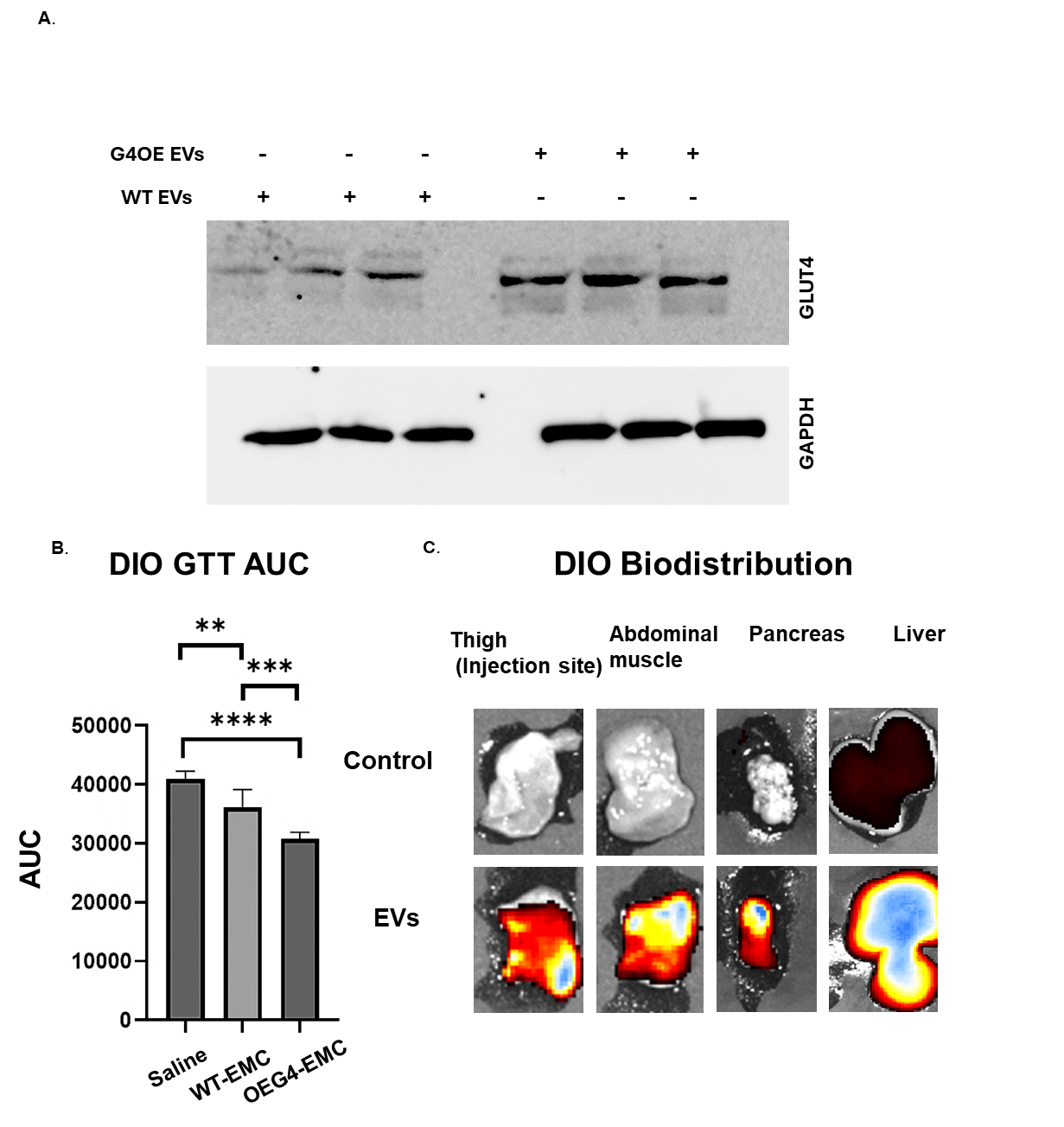

**Figure S2:** **In vivo GTT and EV biodistribution in DIO mice.**

A) Western blot to determine GLUT4 levels in WT-EMC incubated with G4OE-EMC-derived EVs or WT-EMC-derived EVs.
B) Area under the curve analysis of GTT measurements in DIO mice showing integrated improvement in glucose response of mice injected with G4OE-EMC derived EVs.
C) Representative IVIS images of DiR labelled EV biodistribution in DIO mice.

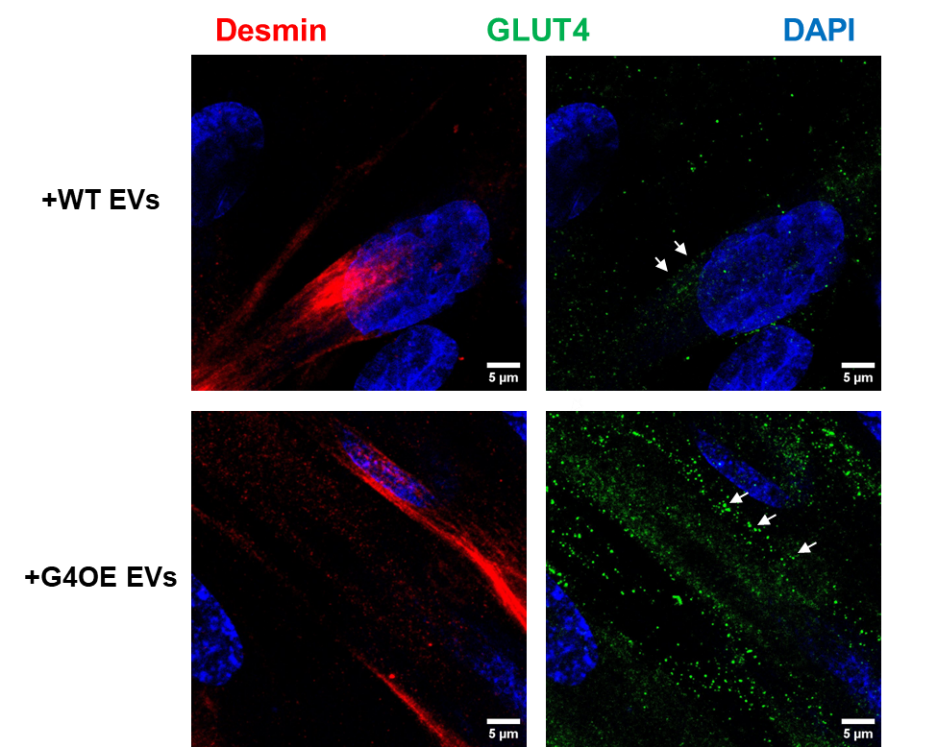

**Figure S3: Enhanced GLUT4 localization following incubation with G4OE-EMC EVs.**

Whole-mount immunostaining of a WT-EMC incubated with G4OE-EMC-derived EVs or WT-EMC-derived EVs and stimulated with insulin, for desmin (Red), GLUT4 (Green) and nucleus (DAPI, Blue), at 63X, as photographed using digital zoom 2X. white arrows indicate GLUT4 localization.
